# ZNF598 responds to mitochondrial stress to abort stalled translation on mitochondrial outer membrane and maintain tissue homeostasis

**DOI:** 10.1101/2022.02.04.479092

**Authors:** Ji Geng, Yu Li, Zhihao Wu, Rani Ohja, Shuangxi Li, Bingwei Lu

**Author notes:** These authors contributed equally (Ji Geng, Yu Li).

## Abstract

Translational control exerts immediate effect on the composition, abundance, and integrity of the proteome. Ribosome-associated quality control (RQC) handles ribosomes stalled at the elongation and termination steps of translation, with ZNF598 in mammals and Hel2 in yeast serving as key sensors of translation stalling and coordinators of downstream resolution of collided ribosomes, termination of stalled translation, and removal of faulty translation products. The physiological regulation of RQC in general and ZNF598 in particular in multicellular settings is underexplored. Here we show that ZNF598 undergoes regulatory K63-linked ubiquitination and its level is upregulated upon mitochondrial stress in mammalian cells and *Drosophila*. Overexpression of ZNF598 protects against mitochondrial stress. In *Drosophila* models of neurodegenerative diseases and patient cells, ZNF598 overexpression aborted stalled translation of mitochondrial outer membrane-associated mRNAs and removed faulty translation products causal of disease. These results shed lights on the regulation of ZNF598 and its important role in mitochondrial homeostasis.

## Introduction

Cells respond to stress stimuli by reconfiguring their transcriptional, translational, and metabolic profiles. Transcriptomics and proteomics studies have revealed that transcript levels and protein abundance do not always match (Vogel and Marcotte, 2012), emphasizing the importance post-transcriptional control of protein output. Compared to transcriptional control, translational control of available mRNAs exerts immediate effect on the composition, abundance, and integrity of the proteome (Hershey et al., 2019), making it particularly important under stress conditions (Advani and Ivanov, 2019). Moreover, protein synthesis or mRNA translation is intrinsically a very energy demanding process, and its regulation is intimately linked to the energy status of the cells (Advani and Ivanov, 2019). Translational control is therefore essential for cellular homeostasis. It critically influences cellular proliferation, growth, and survival, and impacts diverse physiological processes, from early development to synaptic plasticity (Costa-Mattioli et al., 2009; Lasko, 2012). Deregulated translational control is profoundly implicated in diseases (Charmpilas et al., 2015; Kapahi, 2010; Robichaud et al., 2019; Skariah and Todd, 2021; Steffen and Dillin, 2016; Tahmasebi et al., 2018).

Although translation is known to be tightly regulated at the rate-limiting initiation step (Sonenberg and Hinnebusch, 2009), the elongation and termination steps are also subject to intricate regulation (Dever and Green, 2012; Schuller and Green, 2018). During translation elongation, ribosome slowdown and stalling can occur for various reasons. Some are functional and serve to facilitate cellular dynamics, such as co-translational protein folding, frameshifting, and subcellular protein targeting. Others are detrimental and can be triggered by damaged mRNAs, mRNA secondary structures, insufficient supply of aminoacyl-tRNAs, or environmental stress (D’Orazio and Green, 2021; Harigaya and Parker, 2010). Ribosome slowdown and stalling can result in ribosome collision (Kim and Zaher, 2021), which is sensed by the cell as a proxy for aberrant translation and triggers ribosome-associated quality control (RQC) (Brandman and Hegde, 2016; Howard and Frost, 2021; Inada, 2020; Joazeiro, 2019; Sitron and Brandman, 2020). Key factors involved in the process are the ubiquitin ligase ZNF598 (Hel2 in yeast) and the 40S subunit protein Rack1 (Asc1 in yeast), which recognize the distinct 40S-40S interface of collided ribosomes and promote ubiquitination of specific 40S proteins (Juszkiewicz and Hegde, 2017; Sundaramoorthy et al., 2017), and the ASC complex that disassembles the leading collided ribosome (Hashimoto et al., 2020; Juszkiewicz et al., 2020). This then triggers a series of downstream quality control events, including ribosome subunit splitting and recycling by ABCE1 (Shao et al., 2013), CAT-tailing modification by NEMF (Tae2 or Rqc2 in yeast) of nascent peptide chains (NPCs) still attached to the 60S subunit (Shen et al., 2015), release of stalled NPCs from the peptidyl-tRNA/60S complex by ANKZF1 (Vms1 in yeast) (Verma et al., 2018; Zurita Rendon et al., 2018), and degradation of stalled NPCs by the Ltn1 E3 ligase-mediated ubiquitination (Brandman et al., 2012; Shao and Hegde, 2014).

Recent studies have highlighted the importance of regulatory ribosomal protein ubiquitination in the RQC process. RQC is initiated by ZNF598 through site-specific mono-ubiquitination of RPS10/eS10 and RPS20/uS10 of ribosomes stalled on aberrant mRNAs (Garzia et al., 2017; Juszkiewicz and Hegde, 2017; Matsuo et al., 2017; Sundaramoorthy et al., 2017), leading to the proposal that the unique 40S-40S interface of collided ribosomes is specifically recognized by ZNF598 such that problematic translation is differentiated from normal translation. Collided ribosomes marked by ZNF598-dependent ubiquitination of RPS10 are disassembled by the conserved ASC-1 complex (ASCC) containing ASCC3, an ATP-dependent helicase (Hashimoto et al., 2020; Juszkiewicz et al., 2020). In addition to RPS10 and Rps20, other 40S subunits such as Rps2 and Rps3 are also mono-ubiquitinated, with ZNF598 and RNF10 having been implicated in these events (Garzia et al., 2021; Juszkiewicz and Hegde, 2017; Sundaramoorthy et al., 2017). In addition, at least in yeast, the small ribosomal protein eS7 is ubiquitinated by the E3 ligase Not4 (Matsuki et al., 2020). Furthermore, regulatory ribosome ubiquitination involved in RQC is also modulated at the deubiquitination step, with several enzymes having been implicated, including USP10 (Meyer et al., 2020), OTUD3 (Garshott et al., 2020), and USP21 (Garshott et al., 2020) that can remove the ubiquitin added by ZNF598.

Despite our growing knowledge of the players and mechanisms involved in RQC, how the RQC process is regulated by cellular signaling pathways is less well understood. Of particular relevance, the RQC factors are present at sub-stoichiometric levels related to the ribosomal proteins. For examples, yeast Hel2 was reported to be present at less than 1% of the abundance of ribosomes (Warner, 1999). This low abundance of RQC factors relative to ribosomes creates a conundrum under stress conditions when more ribosome collisions have to be dealt with. For example, amino acid deprivation, which leads to depletion of aminoacylated tRNAs, and alkylation and oxidation stresses, which globally damage RNA, are known to increase levels of ribosome collisions (Kim and Zaher, 2021). Although it has been shown that when the RQC pathway is overwhelmed under excess stress and ribosome collision, integrated stress response (ISR) (Meydan and Guydosh, 2020; Yan and Zaher, 2021) and ribotoxic stress response (RSR) pathways (Vind et al., 2020; Wu et al., 2020) can be activated, leading to cell cycle arrest and apoptosis, how the RQC machinery responds to stress under physiological conditions has not been examined.

Here we examined the regulation of ZNF598 under stress in mammalian cells and *in vivo* in *Drosophila*. We found that ZNF598 undergoes regulatory K63-linked ubiquitination and its level is upregulated upon mitochondrial stress. Overexpression of ZNF598 protects against mitochondrial stress in cultured mammalian cells, *Drosophila* models of neurodegenerative disease, and patient cells, by aborting stalled translation of mRNAs associated with mitochondrial outer membrane and removing faulty translation products causal of disease. These results shed new lights on the regulation and function of ZNF598 in mitochondrial homeostasis.

## Results

### Increased ZNF598 expression upon mitochondrial stress in cell culture

To understand how cellular stress might impinge on the RQC process, we examined the expression of ZNF598 under various stress condition. We found that upon serum starvation (EBSS), translational inhibition and autophagy induction with the mTOR inhibitor Torin1, or mitochondrial depolarization with CCCP, only CCCP caused a significant increase of ZNF598 protein level in HeLa cells (Figure 1A, and Figure 1 source data). Treatment of cells with the complex-I inhibitor rotenone also caused an increase of ZNF598, though to a lesser extent than observed in CCCP treated cells (Figure 1B, and Figure 1 source data). The expression level of Rack1, another sensor of ribosome stalling and collision (Sundaramoorthy et al., 2017), was not changed by CCCP treatment (Figure 1B, and Figure 1 source data). Immunostaining of endogenous ZNF598 confirmed CCCP-induced increase of ZNF598 protein level. Intriguingly, the major portion of CCCP-induced ZNF598 was present in the nucleus, with modestly enriched colocalization with mitochondria (Figure 1-figure supplement 1A). The nuclear localization may be related to the known function of ZNF598 homolog Hel2 in yeast to regulate histone homeostasis (Singh et al., 2012), although so far there is no well-defined nuclear function of ZNF598 in mammalian cells. Similar effects were found in cells subjected to a different mitochondrial stress induced by combined inhibition of the electron transport chain with the complex-III inhibitor antimycin A and the complex V inhibitor oligomycin (Figure 1-figure supplement 1A), suggesting that the observed CCCP effect is attributable to mitochondrial stress, not some idiosyncratic effects of CCCP.

**Figure 1.**
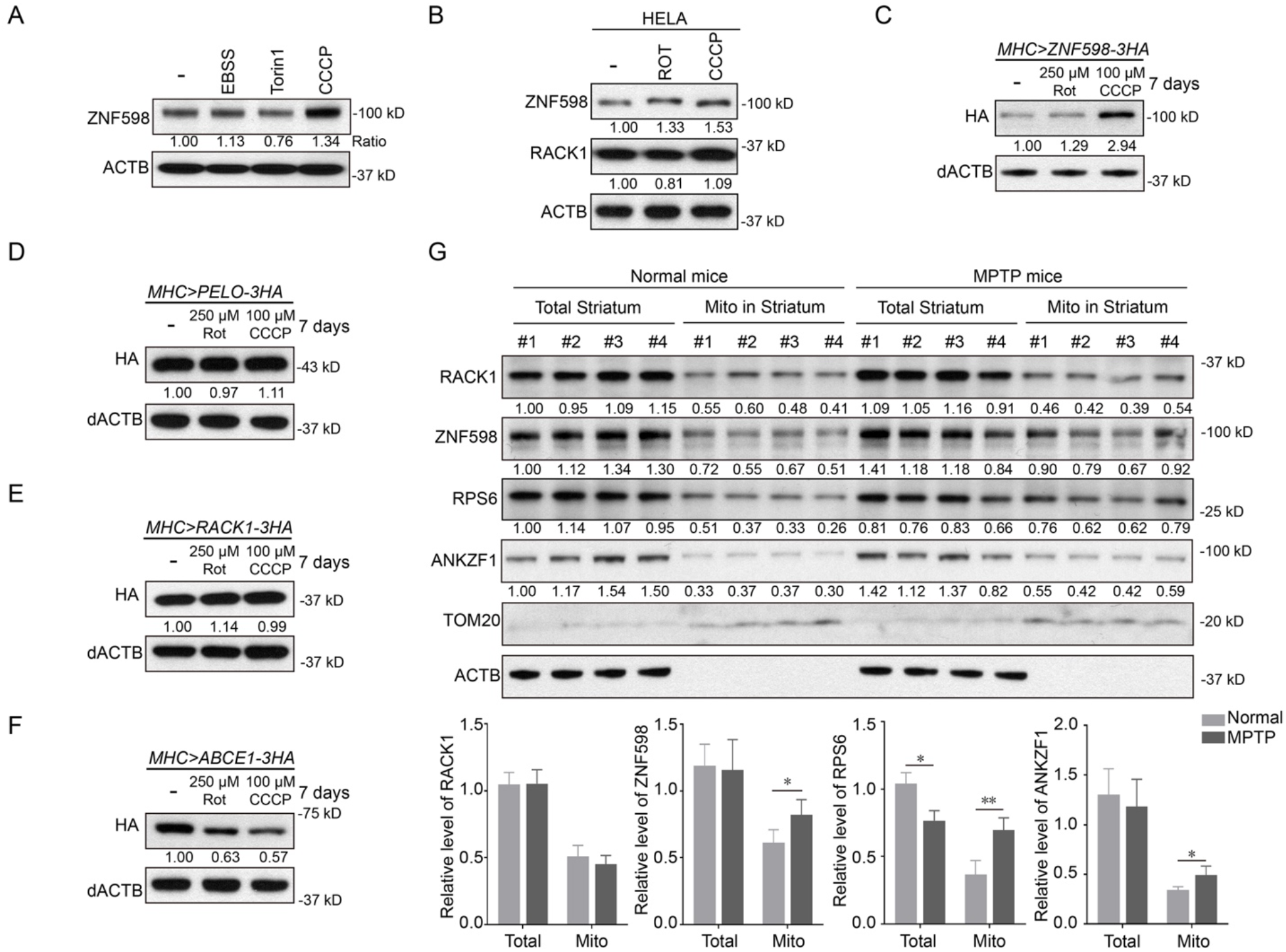
ZNF598 is upregulated under mitochondrial stress conditions *in vitro* and *in vivo*. (**A**) Western blot analysis of ZNF598 protein in HeLa cells treated with EBSS (16 h), Torin1 (0.5 μM, 24 h) and CCCP (10 μM, 24 h). Values under the blot show normalized protein levels relative to untreated sample in this figure and all subsequent ones. (**B**) Western blot analysis of ZNF598 protein in HeLa cells treated with rotenone (5 μM, 24 h) or CCCP (10 μM, 24 h). (**C-F**) Western blot analysis of HA-tagged ZNF598 (**C**), Pelo (**D**), Rack1 (**E**), and ABCE1 (**F**) proteins in the muscle tissue of flies treated with rotenone (250 μM) or CCCP (100 μM) for 7 days. (**G**) Western blot analysis of the indicated proteins in total and mitochondrial fractions isolated from striatum tissues of normal and MPTP-indued mice. Bar graph shows data quantification (n=4). *, P < 0.05 and **, P < 0.01 compared to control group. All data are representative of at least 3 independent experiments. **Figure 1 source data**: Western blots for ZNF598, Rack1, Actin, HA-ZNF598, HA-Pelo, HA-ABCE1, RPS6, ANKZF1, Tom20.

### Increased ZNF598 expression upon mitochondrial stress *in vivo*

We next tested whether the regulation of ZNF598 by mitochondrial stress holds true *in vivo*. For this purpose, we used *Drosophila* as our model system. Due to the lack of antibodies recognizing endogenous fly ZNF598 protein, we made use of a transgene expressing 3xHA-tagged ZNF598. We also expressed 3xHA-tagged Rack1, Pelo, and ABCE1 proteins. Transgenes were expressed in the fly muscle using the *Mhc-gal4* driver and the *UAS-Gal4* system. Transgenic flies were fed for 7 days with fly food containing 250 μM rotenone or 100 μM CCCP. Western blot analysis of protein extracts prepared with dissected thoracis showed that ZNF598-3xHA was significantly increased upon CCCP treatment and moderately increased by rotenone treatment (Figure 1C, and Figure 1 source data). In contrast, Pelo or Rack1 protein levels showed no obvious change (Figure 1D, E, and Figure 1 source data). Consistent with previous findings (Wu et al., 2018), both rotenone and CCCP treatments led to a significant decrease of ABCE1 level (Figure 1F, and Figure 1 source data). In transgenic flies with RNAi-mediated knockdown of PINK1, which causes mitochondrial dysfunction and stress (Yang et al., 2006), ZNF598 level was also increased (Figure 1-figure supplement 1B, and Figure 1-figure supplement source data), consistent with ZNF598 being regulated by mitochondrial stress.

We also tested if ZNF598 protein level responds to mitochondrial stress in a mammalian setting. We used the well-established mitochondrial toxin MPTP (1-methyl-4-phenyl-1,2,3,6-tetrahydropyridine) to induce mitochondrial stress in the mouse brain (Tieu, 2011). MPTP is a prodrug to the neurotoxin MPP^+^, which can be selectively taken up by dopaminergic neurons where it inhibits complex-I activity. We found that compared to Rack1, the level of ZNF598 in the mitochondrial fraction of striatum was moderately increased after MPTP treatment (Figure 1G, and Figure 1 source data). The level of the cytosolic ribosomal protein RPS6 was also increased in the striatum mitochondrial fraction (Figure 1G, and Figure 1 source data), suggesting possible accumulation of cytosolic ribosomes on the mitochondria surface upon MPP^+^-induced mitochondrial stress. ANKZF1 level was also increased in the striatum mitochondrial fraction after MPTP treatment (Figure 1G, and Figure 1 source data). It is worth noting that since MPP+ selectively accumulates in DA neurons and our western blot analysis was performed on whole striatum tissue, the observed changes in ZNF598 and ANKZF1 levels upon MPTP treatment were underestimates of their actual changes in the brain cells actually experiencing MPTP-induced mitochondrial damage.

### ZNF598 protects cells against mitochondrial stress and other stress stimuli

To examine ZNF598 function in mammalian cells, we used CRISPER-Cas9 to knockout its expression. The knockout efficiency was confirmed by Western blot analysis (Figure 2-figure supplement 2A, and Figure 2-figure supplement source data 2). Next, we challenged control and ZNF598 KO cells with various stressors, including the mitochondrial stressors rotenone and CCCP, the ER stressor thapsigargin, and the lysosomal stressor bafilomycin A1. ZNF598 KO cells were more sensitive to these stressors as measured with the CCK8 cell viability assay (Figure 2A, and Figure 2 source data 1), suggesting that ZNF598 normally protects cells against the stress stimuli induced by these agents.

**Figure 2.**
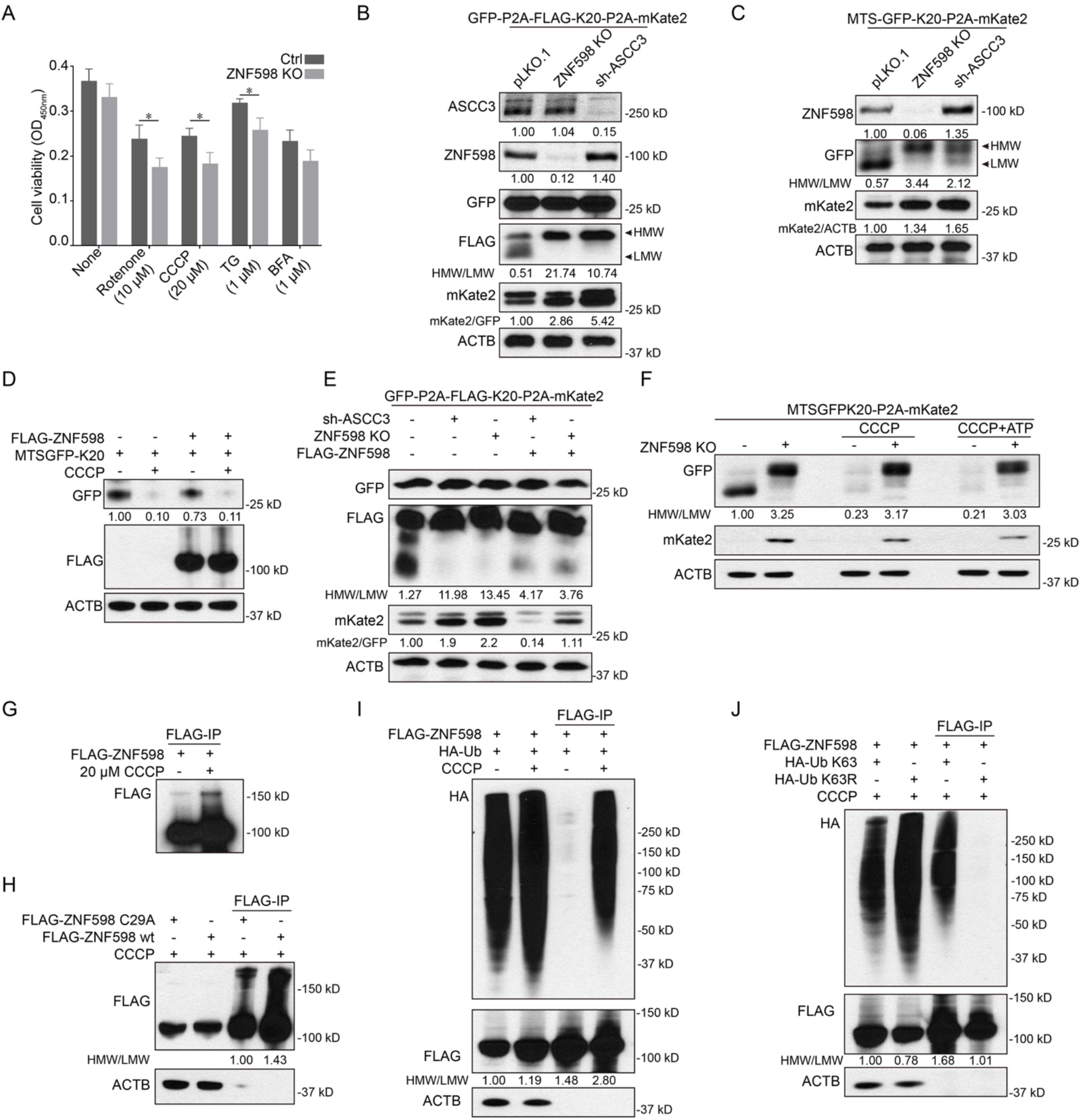
ZNF598 undergoes regulatory K63-linked ubiquitination under mitochondrial stress. (**A**) Cell viability assay of control and ZNF598 KO cells treated with rotenone, CCCP, TG and BFA for 24 h. n=5, *, P < 0.05 and **, P < 0.01 compared with control. (**B**) Immunoblots of GFP-P2A-Flag-K20-P2A-mKate2 stall reporter expression in control (pLK0.1), ZNF598 KO, and ASCC3 KD cells. For anti-Flag blot, the ratio of high MW (HMW) to low MW (LMW) Flag signals corresponding to full-length Flag-K20 and arrested Flag-K20, respectively, was shown. (**C**) Immunoblots of MTS-GFP-K20-P2A-mKate 2 stall reporter expression in control (pLK0.1) ZNF598 KO, and ASCC3 KD cells. For anti-GFP blot, the ratio of HMW to LMW GFP signals corresponding to full-length GFP-K20 and arrested GFP-K20, respectively, was shown. (**D**) Effect of Flag-ZNF598 overexpression on MTS-GFP-K20 stall reporter expression in HeLa cells with or without CCCP treatment (6 hr). (**E**) Immunoblots showing effect of Flag-ZNF598 overexpression on GFP-P2A-Flag-K20-P2A-mKate2 stall reporter expression in control (pLK0.1), ZNF598 KO, and ASCC3 KD cells. (**F**) Effect of ZNF598 KO on MTS-GFP-K20-P2A-mKate 2 stall reporter expression in HeLa cell with or without CCCP treatment or with CCCP + ATP treatment. (**G**) Immunoblots showing effect of CCCP treatment on ZNF598 protein abundance and modification. Immunoprecipitated Flag-ZNF598 protein was analyzed. (**H**) Immunoblots showing effect of CCCP treatment on ZNF598-wt and ZNF598-C29A protein abundance and modification. (**I**) Immunoblots showing effect of CCCP treatment on ZNF598 ubiquitination in cells co-transfected with Flag-ZNF598 and HA-Ub. The signals in the Flag-IP represent ubiquitinated ZNF598 and maybe some ubiquitinated proteins tightly associated with ZNF598. (**J**) Immunoblots showing effect of CCCP treatment on ZNF598 ubiquitination in cells co-transfected with Flag-ZNF598 and HA-Ub-K63 or HA-Ub-K63R. All data are representative of at least 3 independent experiments. **Figure 2 source data 1**: Quantification of cell viability under various stress conditions. **Figure 2 source data 2:** Western blots of ASCC3, ZNF598, GFP, Flag, HA, mKate2, Actin,

We next tested the effect of ZNF598 on stress response *in vivo*. Transgenic flies overexpressing ZNF598 or with ZNF598 knockdown by RNAi were subjected to various stress conditions, including mitochondrial stress, ER stress and starvation. We found that ZNF598 overexpression significantly rescued CCCP- and starvation-induced ATP reduction (Figure 2-figure supplement 2B, and Figure 2-figure supplement source data 1), supporting that ZNF598 helps maintain mitochondrial function during cellular stress.

### ZNF598 regulates quality control of stalled translation of mRNAs encoding mitochondria-targeted proteins

We next tested the mechanism of ZNF598 function in stress response. Using a GFP-P2A-Flag-K20-P2A-mKate2 reporter, in which the GFP, Flag, and mKate2 reporters are used to monitor overall mRNA translation, translational stalling, and readthrough of the stalling site, respectively, and with the self-cleaving P2A peptide sequence allowing each reporter to be independent markers of translation, we assessed the effect of silencing ZNF598 on translation. ZNF598 KO resulted in the removal of arrested translation at the Flag-K20 stall site, allowing increased synthesis of fulllength Flag-K20 and the downstream mKate2 reporter (Figure 2B, and Figure 2 source data 2). GFP expression was not significantly affected. Knocking down ASCC3, a helicase in the ASC-1 complex that selectively dissociates the leading ribosome of a collision, allowing trailing ribosome to continue translation, had similar effect (Figure 2B, and Figure 2 source data 2). These results suggest that in the absence of sufficient ZNF598 or ASCC3 activity, the ribosomes may ignore the stall signal, or that the stalled ribosomes may eventually bypass the stall site if they are not disassembled.

To test whether ZNF598 also regulates the quality control of the translation of mRNAs encoding nuclear-encoded, mitochondrial-targeted proteins, we made use of a mitochondrial-targeted GFP reporter containing the K20 stall signal (MTS-GFP-K20-P2A-mKate2). Co-translational import of mitochondrial proteins plays a critical role in cellular homeostasis (Mohanraj et al., 2020). Given the high frequency of co-translational import of nuclear-encoded mitochondrial proteins as suggested by recent studies (Williams et al., 2014), we expected that at least some of the translation of this reporter would occur on mitochondrial outer membrane. As with the cytosolically localized GFP-P2A-Flag-K20-P2A-mKate2 reporter, the MTS-GFP-K20-P2A-mKate reporter was also regulated by ZNF598 and ASCC3, with the knockout of ZNF598 or knockdown of ASCC3 leading to increased synthesis of full-length GFP-K20 and mKate2 (Figure 2C, and Figure 2 source data 2). On the other hand, overexpression of ZNF598 modestly inhibited MTS-GFP-K20 expression (Figure 2D, and Figure 2 source data 2).

### ZNF598 is upregulated in ASCC3-deficient cells and its overexpression compensates for the effect of ASCC3 loss on translation stalling

We noticed that inhibition of ASCC3 modestly increased ZNF598 protein level, whereas ASCC3 protein levels was not changed in ZNF598 KO cell (Figure 2B, C, and Figure 2 source data 2), suggesting that ASCC3 may negatively regulate ZNF598 abundance, or that the ZNF598 upregulation is a compensatory response to the deficit of ASCC3. We further tested the functional relationship between ZNF598 and ASCC3. We found that overexpression of ZNF598 could partially block ASCC3 RNAi-induced ribosome readthrough of the K(AAA)20 reporter (Figure 2E, and Figure 2 source data 2). Thus, overexpression of ZNF598 may compensate for the effect of loss of ASCC3 on translation stalling.

### Mitochondrial stress activates ZNF598-mediated resolution of stalled translation

We further examined the effect of CCCP-induced mitochondrial stress on MTS-GFP-K20-P2A-mKate2 reporter expression. We found that CCCP treatment resulted in reduced expression of GFP-K20, and mKate 2 was undetectable as in untreated cells (Figure 2D, F, and Figure 2 source data 2). This CCCP effect was unlikely caused by mitochondrial energy deficit as it was not affected by the supplementation of ATP (Figure 2F, and Figure 2 source data 2). The effect was likely due to increased ZNF598 expression as described earlier, and abortive termination of reporter translation by ZNF598. Indeed, knockdown of ZNF598 effectively blocked the CCCP effect, resulting in removal of arrested GFP-K20 and efficient translation of both full-length GFP-K20 and mKate2 (Figure 2F, and Figure 2 source data 2).

The cyclic GMP-AMP synthase-stimulator of interferon genes (cGAS-STING) pathway senses cytosolic DNA and induces interferon signaling to activate the innate immune system. Translation stress and collided ribosomes were recently shown to serve as coactivators of cGAS (Wan et al., 2021). Ribosome collision leads to cytosolic localization of cGAS, which preferentially interacts with collided ribosomes, and this ribosome association stimulates cGAS activity. Using cytosolic cGAS localization as a proxy of its activation in response to ribosome stalling (Wan et al., 2021), we found that anisomycin-induced ribosome collision and ZNF598 KO both increased cytosolic cGAS and decreased nuclear cGAS signals, as previously reported (Wan et al., 2021). However, while CCCP-treatment antagonized the anisomycin effect on nuclear vs. cytoplasmic cGAS localization, it failed to do so in ZNF598 KO cells (Figure 2-figure supplement 2C, and Figure 2-figure supplement source data 1), suggesting that ZNF598 mediates the effect of CCCP in resolving collided ribosomes and aborting stalled translation.

### Regulatory K63-linked ubiquitination of ZNF598 upon mitochondrial stress

We noticed that under CCCP treatment condition, not only ZNF598 protein level was increased, but also some high molecular weight (HMW) species of ZNF598, presumably ubiquitinated ZNF598, was observed (Figure 2G, and Figure 2 source data 2). ZNF598 level was slightly increased after MG132 treatment, suggesting that the proteasome plays a modest role in ZNF598 regulation (Figure 2-figure supplement 2D, and Figure 2-figure supplement source data 2). To test if the HMW species of ZNF598 reflected autoubiquitination, a mechanism of E3 ligase selfregulation (de Bie and Ciechanover, 2011), we tested an E3-activity deficient mutant of ZNF598 (ZNF598-C29A). We found that the HMW species of ZNF598 was decreased but not obliterated for ZNF598-C29A after CCCP treatment (Figure 2H, and Figure 2 source data 2), suggesting that CCCP-induced ZNF598 ubiquitination may involve autoubiquitination as well as ubiquitination by unidentified E3 ligase(s).

To analyze CCCP-induced ubiquitination of ZNF598 further, we co-transfected Flag-tagged ZNF598 and HA-tagged ubiquitin and found that ZNF598 immunoprecipitated with anti-Flag was heavily ubiquitinated under CCCP treatment condition (Figure 2I, and Figure 2 source data 2). Moreover, we co-transfected ZNF598 and mutant forms of ubiquitin and found that mutant Ub with K63 as the only available linkage (Ub-K63) recapitulated the effect of CCCP on ZNF598 ubiquitination, whereas mutant Ub with only K63 linkage blocked (Ub-K63R) prevented the ability of CCCP to induce formation of the ubiquitinated ZNF598 species (Figure 2J, and Figure 2 source data 2), suggesting that CCCP-induced mitochondrial stress promotes K63-linked ubiquitination of ZNF598. Moreover, the association of this K63 ubiquitination with higher protein levels and higher activity of ZNF598 suggests that this ubiquitination event plays a regulatory and signaling function.

### Regulation by ZNF598 of stalled translation of mitochondrial outer membrane-associated *C-I30* mRNA in *PINK1* mutant

To test the *in vivo* function of ZNF598 in translational quality control of physiological substrates, we used the translation of mitochondrial complex-I subunit *C-I30* mRNA as a model. Previous studies showed that the translation of *C-I30* mRNA occurs on mitochondrial outer membrane (Gehrke et al., 2015), which is sensitive to mitochondrial stress or the loss-of-function of the mitochondrial quality control factor PINK1, mutations in which are associated with Parkinson’s disease (PD) (Valente et al., 2004). Mitochondrial dysfunction causes ribosome stalling at the canonical stop codon site due to the deficiency of translational termination and ribosome recycling activities and resulted in the formation of CAT-tailed C-I30-u species, which are toxic and contribute to PINK1 pathogenesis (Wu et al., 2019). First, we tested whether ZNF598 and other RQC factors are localized to mitochondria *in vivo*. We found that ZNF598 and other RQC factors Rack1, Pelo, Vms1/ANKZF1, and VCP showed various degrees of localization to mitochondria in the DA neurons of adult fly brain (Figure 3A, Figure 3-figure supplement 3A). Next, we tested the effect of overexpression of ZNF598 and the RQC factors on C-I30-u formation. We found that ZNF598 had the strongest effect in eliminating C-I30-u (Figure 3-figure supplement 3B, and Figure 3-figure supplement source data 2). Overexpression of the other RQC factors also reduced C-I30-u level by varying degrees. For example, overexpression of HA-Rack1 had no obvious effect on C-I30-u level (Figure 3B, and Figure 3 source data 2). Importantly, compared to control flies, *PINK1* mutant flies accumulated several fold higher levels of HA-ZNF598 (Figure 3B, and Figure 3 source data 2). These results demonstrates that ZNF598 responds to mitochondrial stress *in vivo* and plays an important role in the quality control of mitochondria-associated ribosome stalling.

**Figure 3.**
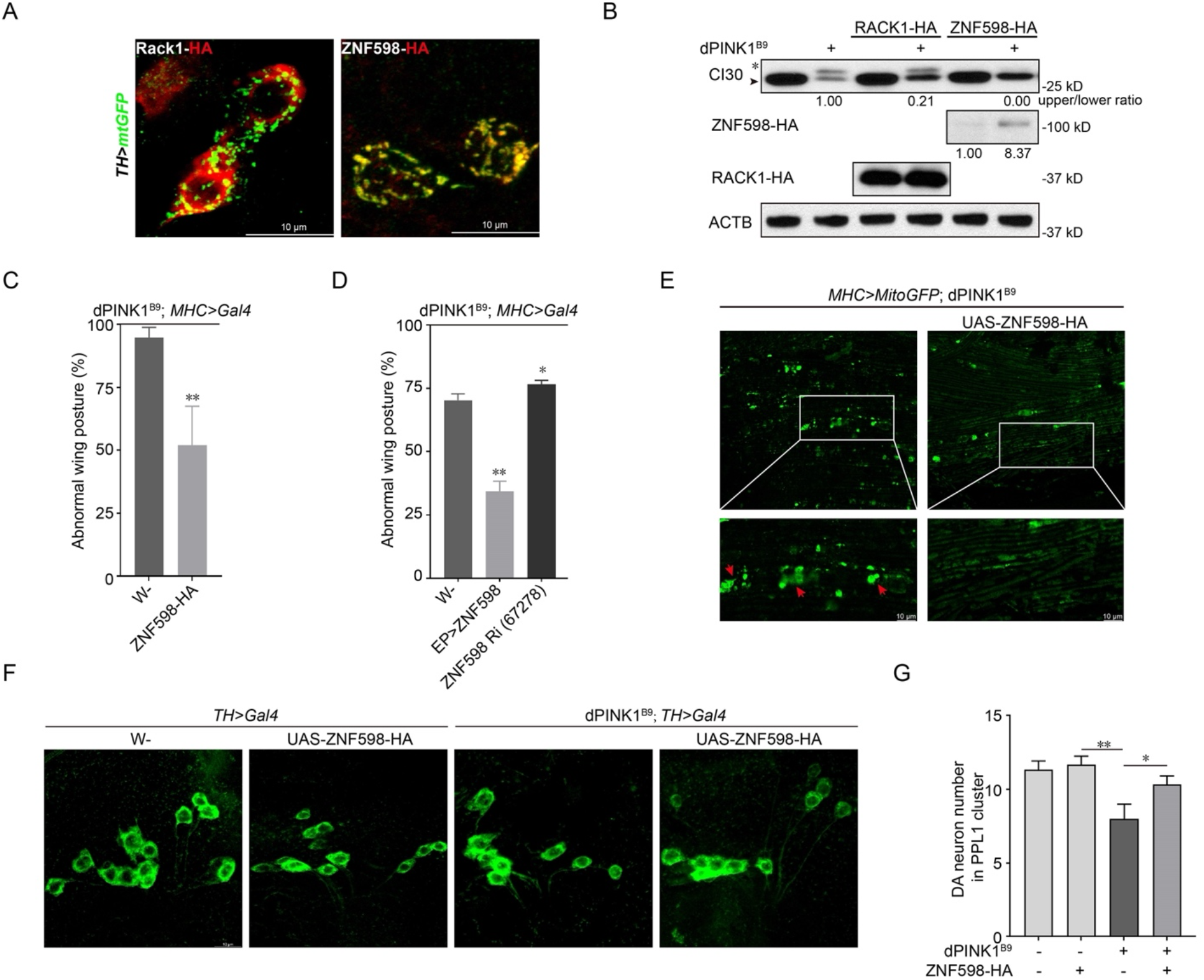
ZNF598 aborts stalled translation of mitochondrial outer membrane-associated *C-I30* mRNA and rescues mitochondrial and neuromuscular defects of *PINK1* mutant flies. (**A**) Localization of HA-tagged early RQC factors ZNF598 and Rack1 to mitochondria of adult DA neurons. DA neuron mitochondria are marked with TH-Gal4-driven mito-GFP expression. (**B**) Western blot analysis showing effect of ZNF598 and Rack1 overexpression on CAT-tailed C-I30-u formation in *PINK1^B9^* fly muscle. (**C, D**) Effect of ZNF598 overexpression (by UAS-ZNF598-HA or ZNF598-EP) or RNAi on abnormal wing posture in *PINK1^B9^* flies. n=3, *, P<0.05, **, P < 0.01 compared with *PINK1^B9^* control flies. 10-20 flies per genotype per assay. (**E**) Effect of ZNF598 overexpression on mitochondrial morphology in *PINK1^B9^* fly muscle. Red arrowheads mark aggregated mitochondria. Mito-GFP labels mitochondria. (**F, G**) Effect of ZNF598 overexpression on DA neuron number in the PPL1 cluster of *PINK1^B9^* adult brains. Bar graph shows data quantification (**G**). n=3, *, P<0.05, **, P < 0.01 compared with *PINK1^B9^* control flies. 5-10 brain samples per genotype per assay. All data are representative of at least 3 independent experiments. **Figure 3 source data 1**: Quantification of wing posture and DA neuron number. **Figure 3 source data 2**: Western blots of C-I30, HA-ZNF598, HA-Rack1, and Actin.

### Overexpression of ZNF598 rescues PD-related phenotypes *in PINK1* mutant

Consistent with the effect of ZNF598 on C-I30-u level, which strictly correlated with disease severity in *PINK1* mutant as shown previously (Wu et al., 2019), overexpression of ZNF598 in the muscle effectively rescued the abnormal wing posture phenotype in *PINK1* mutant (Figure 3C, D, and Figure 3 source data 1), which was caused by mitochondrial dysfunction-induced indirect flight muscle degeneration (Yang et al., 2006). This was correlated with significant rescue of mitochondrial morphology as shown by the removal of defective, swollen mitochondria (Figure 3E). We also examined the effect of ZNF598 overexpression on the PD-related DA neuron loss phenotype in *PINK1* mutant. ZNF598 effectively rescued the neuronal loss in the PPL1 cluster DA neurons in the *PINK1* mutant adult brain (Figure 3F, G, and Figure 3 source data 1), concomitant with the rescue of mitochondrial morphology (Figure 3-figure supplement 3C, and Figure 3-figure supplement source data 1). These data support the important role of ZNF598 in promoting mitochondrial and tissue homeostasis in a PD model.

### ZNF598 regulates the quality control of C9ALS-associated poly(GR) translation stalled on mitochondrial surface in *Drosophila*

We sought to further test the *in vivo* effect of ZNF598 on mitochondria-associated translational quality control. Expansion of G4C2 repeats in the *C9orf72* gene causes amyotrophic lateral sclerosis with frontotemporal dementia (C9-ALS/FTD), one of the most common forms of ALS. The dipeptides translated from G4C2 repeat transcripts by unconventional translation, especially the arginine-containing poly(GR) and poly(PR), are considered disease-relevant toxic species (Gendron and Petrucelli, 2018; Taylor et al., 2016; Yuva-Aydemir et al., 2018). We previously found that the translation of poly(GR) can occur on mitochondrial surface, presumably due to GR repeats mimicking mitochondrial-targeting signal, and ribosomes translating poly(GR) are frequently stalled, seemingly caused by positively-charged Arg residues interacting with negatively-charged residues lining the ribosome exit channel (Li et al., 2020b). Stalled translation of poly(GR) triggers CAT-tailing-like C-terminal extensions, which promote poly(GR) aggregation and toxicity (Li et al., 2020b). We found that overexpression of ZNF598 dramatically reduced the levels of both the CAT-tailed and non-CAT-tailed poly(GR) protein species, with the CAT-tailed species running as a smear slightly above the non-CAT-tailed species (Figure 4A, and Figure 4 source data 2). The reduction of poly(GR) protein expression by ZNF598 was also evident in immunostaining of muscle tissues (Figure 4-figure supplement 4A). This was correlated with the rescue of the neuromuscular toxicity of poly(GR) as measured with wing posture defects in *Mhc*>*GR80* flies (Figure 4B, and Figure 4 source data 1). RNAi of ZNF598 had opposite effects (Figure 4C, D, and Figure 4 source data 1 and Figure 4 source data 2). The reduction of poly(GR) protein expression by ZNF598 correlated with its rescue of the mitochondrial morphology defect of *Mhc*>*GR80* fly muscle (Figure 4E). These data support the important role of ZNF598 in promoting mitochondrial and tissue homeostasis in an ALS model.

**Figure 4.**
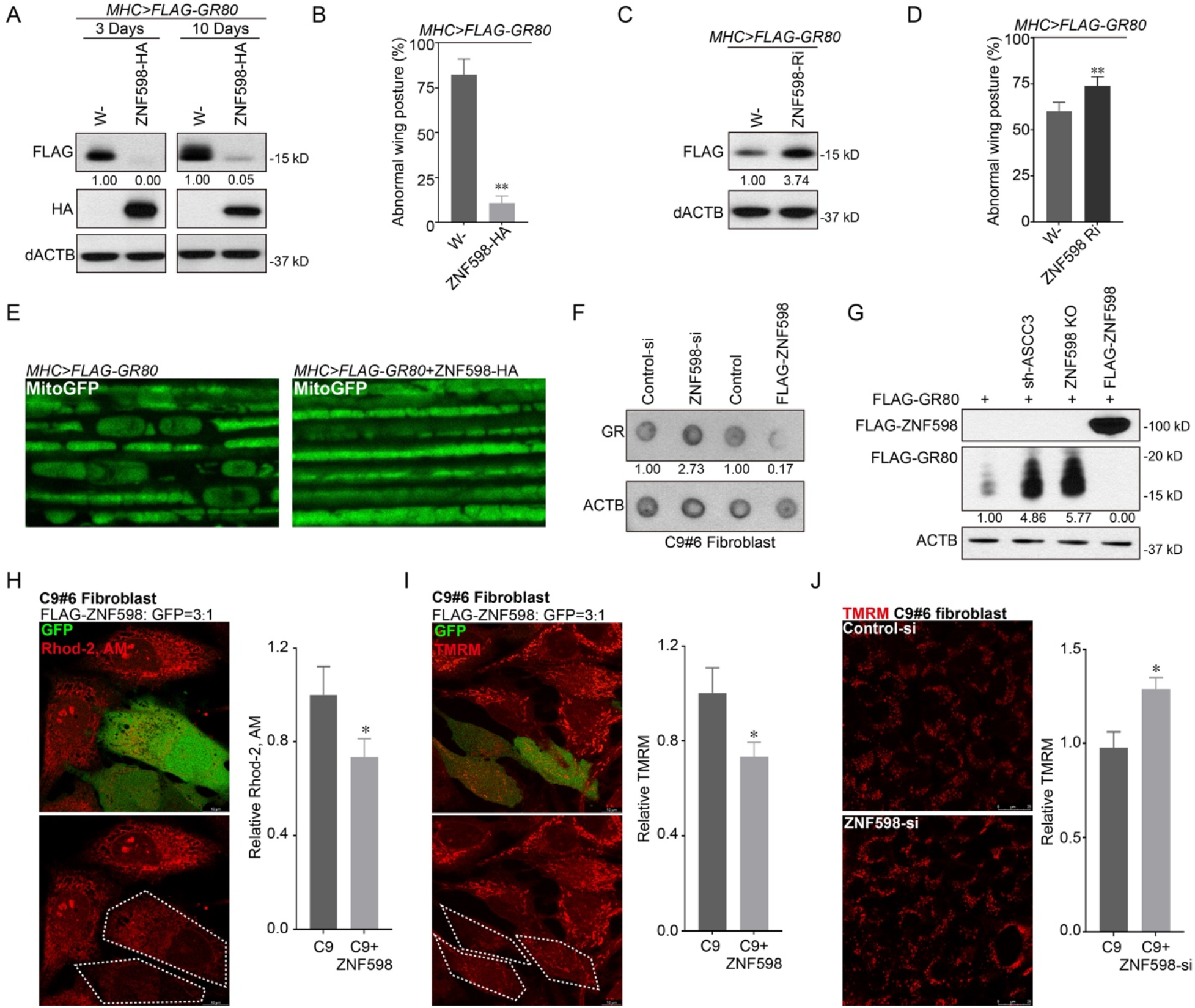
ZNF598 regulates the quality control of stalled translation of C9ALS-associated poly(GR) and rescues poly(GR) induced toxicity in *Drosophila* and patient cells. (**A**) Immunoblots showing effect of ZNF598-HA overexpression on the level of Flag-tagged GR80 expressed in fly muscle. (**B**) Effect of ZNF598-HA overexpression on wing posture in *Mhc*>*GR80* flies. n=5, **, P < 0.01. 10-20 male flies per genotype per assay. (**C**) Immunoblots showing effect of ZNF598 RNAi on the level of Flag-tagged GR80 in *Mhc*>*GR80* flies. (**D**) Effect of ZNF598-RNAi on wing posture in *Mhc*>*GR80* flies. n=5, **, P < 0.01. 10-20 male flies per genotype per assay. (**E**) Effect of ZNF598 overexpression on mitochondrial morphology in *Mhc*>*GR80* fly muscle, which exhibits vacuolated and disconnected mitochondria. (**F**) Dot blots showing effect of ZNF598 overexpression or RNAi on poly(GR) protein level in C9-ALS patient fibroblasts. (**G**) Immunoblots showing effect of ASCC3 or ZNF598 silencing, or ZNF598 overexpression on Flag-GR80 protein expression in HeLa cells. (**H, I**) Effect of ZNF598 on mitochondrial calcium stained by Rhod-2AM (**H**) and MMP stained by TMRM (**I**) in C9-ALS fibroblasts. GFP was co-transfected with ZNF598 at 1:3 ratio so that virtually all ZNF598 transfected cells would be marked with GFP. Bar graphs show data quantification. n=5, *, P < 0.05. (**J**) Effect of ZNF598 RNAi on MMP stained by TMRM in C9-ALS fibroblasts. Bar graph shows data quantification. n=5, *, P < 0.05. All data are representative of at least 3 independent experiments. **Figure 4 source data 1**: Quantification of wing posture, mitochondrial calcium, and MMP. **Figure 4 source data 2**: Western blots of Flag-GR80, ZNF598-HA, Actin, Flag-ZNF598, and dot blots of poly(GR) and Actin.

### ZNF598 regulation of poly(GR) expression and toxicity in C9ALS patient fibroblasts

We next tested ZNF598 function in regulating poly(GR) translation in a more physiologically relevant disease setting using C9-ALS/FTD patient fibroblasts. We found that overexpression of ZNF598 resulted in decreased poly(GR) expression, whereas ZNF598 RNAi had opposite effect in C9-ALS/FTD patient fibroblasts (Figure 4F, and Figure 4 source data 2). Consistent with this result, GR80 protein expression in HeLa cells was significantly increased when ZNF598 or ASCC3 was silenced, but dramatically decreased when ZNF598 was overexpressed (Figure 4G, and Figure 4 source data 2). Moreover, aggregates of GR80 were observed in the nuclei in ZNF598 or ASCC3 silenced cells (Figure 4-figure supplement 4B). Furthermore, overexpression of ZNF598 totally block the effect of ASCC3 silencing on GR80 expression (Figure 4-figure supplement 4C, and Figure 4-figure supplement source data 2), again supporting the notion that overexpression of ZNF598 may compensate for the effect of loss of ASCC3 on translation stalling.

As reported previously (Li et al., 2020a), C9-ALS/FTD patient fibroblasts exhibited abnormal mitochondrial morphology and elevated mitochondrial membrane potential (MMP) and mitochondrial Ca^2+^ compared to cells from control subjects. The increased MMP in C9-ALS fibroblasts is likely due to the alteration and tightening of cristae junctions caused by poly(GR), thereby impairing mitochondrial ion homeostasis (Li et al., 2020a). We found that overexpression of ZNF598 restored mitochondrial morphology (Figure 4-figure supplement 4D, and Figure 4-figure supplement source data 1), and reduced mito-Ca^2+^ level (Figure 4H, and Figure 4 source data 1) and MMP (Figure 4I, and Figure 4 source data 1) in C9-ALS/FTD patient fibroblasts. On the other hand, ZNF598 RNAi resulted in increased MMP (Figure 4J, and Figure 4 source data 1). These result strongly support the relevance of ZNF598 regulation of stalled poly(GR) translation to mitochondrial homeostasis and C9-ALS/FTD pathogenesis.

## Discussion

The translational machinery is intimately linked to environmental conditions, making ribosomes excellent candidates for sensors of the cellular state and platforms for various signaling pathways that respond to cellular changes. In particular, ribosome collision frequency is considered a rheostat used by the cell to select the most appropriate response to problems encountered during translation (Kim and Zaher, 2021). Under normal conditions or when stresses are manageable, cells may use the translation factor eIF5A to handle naturally occurring stalls (Han et al., 2020) or the RQC pathway to resolve infrequent collisions that result from aberrant mRNAs. This typically result in resumption of translation. Under these conditions, maintaining RQC factors such as ZNF598 at sub-stoichiometry relative to ribosomes may be advantages to cells, as too much ZNF598 activity may cause abortive translation of stalls that serve physiological purposes. The situation becomes more complicated with more severe stress conditions when ribosome collision increases. Given the low abundance of RQC factors relative to ribosomes, it is conceivable that the RQC pathway will be overwhelmed under these conditions, necessitating global stress response or cell death (Kim and Zaher, 2021). Our results indicate that before cells succumb to stress, there is an orchestrated RQC response that upregulates the activity and abundance of ZNF598. This upregulation on demand helps alleviate the substoichiometry issue of ZNF598 and is important for maintaining mitochondrial and tissue homeostasis under stress. Our results resonate with the emerging concept that signaling on collided ribosomes has consequences beyond that of just ribosome rescue and mRNA quality control to include triggering of global stress responses (Meydan and Guydosh, 2020; Yan and Zaher, 2021), including the cGAS-STING innate immune response (Wan et al., 2021), and cell fate decisions (Vind et al., 2020; Wu et al., 2020). We hypothesize that this ZNF598 regulation by upstream stress signaling pathways may mechanistically link proteostasis, mitochondrial homeostasis and innate immune response, failures of which are hallmarks of neurodegenerative diseases.

Our results show that ZNF598 responds to mitochondrial stress, and that its upregulation promotes the quality control of stalled cytoplasmic ribosomes associated with mitochondrial surface and the clearance of faulty translation products causal of disease in animal models of PD and ALS. The importance of ZNF598 upregulation to mitochondrial and tissue homeostasis is consistent with previous studies implicating the crosstalk between cytosolic translation and mitochondrial function (Mohanraj et al., 2020), and the particular involvement of the RQC pathway in maintaining mitochondrial homeostasis (Izawa et al., 2017; Wu et al., 2019). The nature of the mitochondrial signal that leads to ZNF598 ubiquitination remains to be determined. There are a few possible candidates. Mitochondrial outer membrane is known to be decorated with cytosolic ribosomes engaging in co-translational import of nuclear encoded mitochondrial proteins or proteins mistargeted to mitochondria, including the C-I30 and poly(GR) proteins studied here in the PD and ALS models, respectively (Gehrke et al., 2015; Li et al., 2020b). Mitochondrial import is known to be sensitive to MMP and other parameters of mitochondrial function. Thus, defects in the co-translational import process may lead to translation stalling and ribosome collision under mitochondrial stress. It is also possible that mitochondrial dysfunction may lead to excessive ROS production, leading to damage of mitochondria-associated mRNA. Oxidizing agents are known to modify the nucleobases of mRNAs, resulting in adducts such as 1-methyladenosine and 8-oxoguanosine, which inhibit tRNA selection by ribosomes and cause their arrest (Kim and Zaher, 2021). In yeast, oxidative stress has been shown cause a decrease in the levels of charged Trp-tRNA and ribosome stalling at Trp codons (Rubio et al., 2021). Whether similar events may occur in metazoans remains to be tested. Other signals derived from mitochondrial stress, such as mitochondrial retrograde signals or mitochondrial unfolded protein response may also impinge on the RQC pathway. Future studies will test these possibilities.

Our results showed that in response to mitochondrial stress, cellular RQC activity in aborting stalled translation was increased in a ZNF598-dependent manner and that ZNF598 became polyubiquitinated in the process, apparently through a combination of autoubiquitination and ubiquitination by other E3 ligase(s). Understanding the effect of ubiquitination on ZNF598 function and biochemical activity requires further studies determining the mitochondrial stress-induced ubiquitination site(s) and testing their functional role through mutagenesis studies. Possible role of other post-translational modifications such as phosphorylation in regulating mitochondrial stress-induced ZNF598 ubiquitination is also worth exploring. Identifying the E3 ligase(s) directly involved in ZNF598 ubiquitination will also be an important future direction. Given the effect of PINK1 in regulating RQC factor recruitment to mitochondrial outer membrane (Wu et al., 2018) and in regulating ZNF598 protein level as reported here, and the effect of ZNF598 overexpression in rescuing *PINK1* mutant phenotypes, genes in the PINK1 pathways are worth considering. In this respect, the E3 ligase Parkin is a good candidate as Parkin is known to regulate mitochondria-associated C-I30 translation (Gehrke et al., 2015) and carry out K63-linked ubiquitination of substrates (Lim et al., 2005). The E3 ligase NOT4 is another potential upstream regulator of ZNF598. NOT4 also responds to mitochondrial stress and promotes the ubiquitination of the RQC factor ABCE1 (Wu et al., 2018). Not4 was originally identified as a conserved component in the CCR4-NOT RNA quality control complex, but its importance in co-translational RQC (Collart and Weiss, 2020), including regulatory ribosomal ubiquitination (Matsuki et al., 2020), is increasingly being recognized. NOT4 therefore represent a potential link between mitochondrial stress-induced RNA damage and RQC. Previous studies also implicated a non-canonical Notch signaling pathway in regulating RQC (Li et al., 2020b). It would be interesting to test its possible role in ZNF598 regulation. Finding the upstream regulators of ZNF598 in response to mitochondrial stress will offer new insight into the regulation of RQC and how defects in this process may contribute to the pathogenesis of neurodegenerative diseases.

## Materials and methods

### *Drosophila* genetics

All *Drosophila* stocks were maintained at 25°C on standard food incubators with a 12 h light:dark cycle. The fly stocks were obtained from the following sources: UAS-mito-GFP (William Saxton, University of California at Santa Cruz, Santa Cruz, CA), *PINK1^B9^* (Jongkeong Chung, Seoul National University, Seoul, Republic of Korea), UAS-FLAG-GR80 (Fenbiao Gao, UMass Medical School); The UAS-Rack1-HA, -Pelo-HA, -ZNF598-HA, -ABCE1-HA, -VCP-HA, - Vms1/ANKZF1-HA flies are purchased from FlyORF. The ZNF598 EP line, and ZNF598 RNAi fly lines are purchased from the Bloomington *Drosophila* Stock Center. Other stocks were generated in our laboratory. Fly culture and crosses were performed according to standard procedures. Adult flies were generally raised at 25°C and with 12/12 hr dark/light cycles. Fly food was prepared with a standard receipt (Water, 17 L; Agar, 93 g; Cornmeal, 1,716 g; Brewer’s yeast extract, 310 g; Sucrose, 517 g; Dextrose, 1033 g).

### Wing posture assays

Around 10-20 male flies were transferred to a clean plastic vial. To assay wing posture, cohorts of flies raised at 25 degrees at the indicated ages were visually for straight, held-up, or droopy wing postures. The number of flies with normal (straight) or abnormal (held-up or droopy) wing postures were counted and quantified as the percentage of the total number of flies.

### Western blot and dot blot assays

Cultured cells and fly thorax muscles were lysed in lysis buffer (50 mM Tris-HCl pH7.4, 150 mM NaCl, 10% glycerol, 1% Triton X-100, 5 mM EDTA) containing protease/phosphatase inhibitor (B14001/B15001, Bimake). After centrifugation at 14,000 × g (4°C) for 15 min and denaturation by boiling in loading buffer, samples containing 30 μg proteins were separated by 12.5% SDS-PAGE and transferred onto PVDF membranes. To detect GR repeats in C9 ALS fibroblast, dot blot was applied as described before (Li et al., 2020a). 10 μg proteins were drop in the PVDF membrane directly. After blocking with 5% nonfat milk at room temperature for 1 h, membranes were incubated with primary antibodies at 4°C overnight. Membranes were rinsed three times with TBST (0.1% Tween20 in TBS) for 10 min each and incubated with HRP-conjugated secondary antibody for 1 hour. After being washed for three times with TBST for 10 minutes each, the membrane was completely immersed with ECL substrate (Perkin Elmer, Beaconsfield, UK) for 1 min. Finally, membrane was exposed to autoradiography film in a darkroom. Intensity of immunoblot bands was quantified using Fiji/ImageJ software. Quantitative analysis of protein levels was determined and is expressed as a ratio to β-actin. Relative quantification was measured as a ratio to control samples.

### MPTP-induced PD mice model study

C57BL/6 mice (male, 10 weeks old, 23-25 g) were purchased from the Jackson Laboratory. Animal welfare and experimental procedures were carried out strictly in accordance with the related ethical regulations of Stanford university. MPTP-PD mice model was conducted as described previously (Geng et al., 2019). Adult male C57Bl/6 mice were administered intraperitoneally with MPTP (dissolved in PBS) in a final concentration of 25 mg/kg daily for five consecutive days and maintained for another 7 days before sacrifice. Striatum was dissected to isolate mitochondria. In brief, striatum tissue was manually homogenized in homogenization buffer (30 mM Tris-HCl pH 7.4, 225 mM mannitol, 75 mM sucrose, 0.5 mM EGTA, protease inhibitor and 0.5% BSA) on ice. Nuclei and unbroken cell debris were pelleted by centrifugation at 700 × g for 5 min. The supernatant was collected and centrifuged at 6300 × g for 10 min at 4 °C, and the pellet is taken as crude mitochondria fraction.

### Immunofluorescence microscopy

For immunostaining of fly thorax and brain tissues, flies were anesthetized using CO_2_ enriched air, and thorax or brain tissues were dissected and fixed in 4% paraformaldehyde containing 0.3% Triton X-100 at 4°C overnight. The thoraces were subdivided into small chunks. The brain was removed from the head cuticle, and carefully removed the surrounding trachea. Samples were permeabilized with 0.5% Triton X-100 in PBS for 45 min before blocking.

The cells were fixed with 4% paraformaldehyde at room temperature for 15 min, permeated with 0.5% Triton X-100 for 15 min, blocked with 5% BSA for 1 h, and incubated with primary antibodies overnight. After washing 3 times with 0.1% Triton X-100 for 10 min, the cells were incubated with secondary antibodies for 1 h at room temperature, washed three times with 0.1% Triton X-100, and finally mixed with mounting medium with DAPI. The images were obtained with a laser confocal microscope Leica SP8 (Leica, Germany). The immunofluorescence intensity and distribution were further analyzed using Fiji/Image J.

### Plasmid construction and transfection

The GFP-P2A-FLAG-K(AAA)20-P2A-mKate2 construct was modified based on GFP-P2A-FLAG-K(AAG)20-P2A-RFP (105689, Addgene). For MTS (Mitochondrial targeting sequence)- GFP-K(AAA)20-P2A-mKate2 construction, the GFP-P2A-FLAG fragment was removed from GFP-P2A-FLAG-K(AAA)20-P2A-mKate2 plasmid, and the GFP fragment and MTS sequence derived from the first 29 AA of cytochrome c oxidase subunit 8A (COX8A) were inserted into the digested construct by using Gibson Assembly HiFi Master Mix (A46627, Invitrogen).

Cells were cultured at 70-80% confluence in 6-well plates before transfection. Lipofectamine 2000 reagent (2.5 μl) was diluted with 100 μl Opti-MEM and gently blended with 100 μl Opti-MEM containing 2.5 μg plasmid. The mixture was placed at room temperature for 5 min and then added to the cell culture medium for 24 h.

### Cell culture conditions

Regular HeLa cells and HEK293T cells (ATCC) were cultured under standard conditions (1x DMEM medium, 5% FBS, 5% CO2, 37°C). C9-ALS patient fibroblasts and matched control fibroblasts were described before (Kramer et al. 2016) and kindly provided by Dr. Aaron Gitler. Hela, HEK 293T, and fibroblast cell transfections were performed by using Lipofectamine 3000 (cat#: L3000015, Invitrogen), and si-RNA knockdown experiments were performed using Lipofectamine RNAiMAX reagent (cat#: 13778150, Invitrogen), according to manufacturer’s instructions.

### Gene knockdown and knockout in cell culture

To reduce ASCC3, shRNA targeting ASCC3 (5-CTACTTCAAAGGCGGTATATA-3) were synthesized and cloned into pLKO.1 vector. pRSV-Rev, pMDL-RRE, pVSV-G were mixed with pLKO.1 at the ratio of 1:1:1:2 and transfected into HEK293T cells for 72 h, the lenti particle in supernatant was concentrated by using PEG-8000. To knockout ZNF598, CRISPR/Cas9 was used. Three sgRNAs targeting ZNF598 (sg1: 5-CCGATGACACGTCGCACCGT-3; sg2: 5-AGGCGGTAGGCCCAAGAAGG-3; sg3: 5-CTACTGCGCCGTGTGCCGCG-3) were synthesized and cloned into CRISPR V2 vector. Viruses were produced in HEK293T cells using psPAX2, pMD2.G and CRISPR V2, and concentrated using PEG-8000, and divided into small aliquot and stored in the −80°C fridge. Cells were plated in 6-well plates in medium with 10% FBS and 2 μg/ml polybrene and infected with 20 μl of virus. The medium was refreshed, and puromycin (2 μg/ml) was added for selection after 48 h.

### Co-IP assay

Cells were homogenized in lysis buffer. After centrifuging at 14,000 g and 4°C for 15 min, the supernatant was immediately transferred to new tubes. Protein concentration was measured using BCA kit. 10 μl M2-anti-FLAG beads were added into lysis buffer and incubated overnight at 4°C on a rocker. Then, the tubes were centrifuged at 500 g for 5 min, the pellet was washed 3 times with precooled PBST, and the beads were boiled in loading buffer. The supernatants were collected and subjected to Western blot analysis.

### CCK8 cell viability assay

The viability of cells was determined using Cell Counting Kit-8 (CCK8) assays (HY-K0301, MCE) with the number of viable cells being evaluated after 2 h in medium containing CCK8. The conversion of the tetrazolium salt WST-8 to formazan was measured at 450 nm using a plate reader.

### Mitotracker Red, Rhod-2AM, and TMRM stainings

For mitotracker staining, cells were washed with PBS and loaded with 500 nM MitoTracker Deep Red (M22425, Invitrogen) for 30 min. Cells were washed with PBS twice to remove the extra dyes. The fluorescent images were captured by microscope Leica SP8, and intensity was measured by Fiji/Image J. For Rhod-2AM staining, cells were washed with PBS and loaded with 5 μM Rhod-2 AM (R1244, Invitrogen) in HBSS without calcium/magnesium for 1 h. Cells were washed with HBSS twice to remove the extra dyes. The fluorescent images were captured by microscope Leica SP8, and intensity was measured by Fiji/Image J. MMP was measured by using Image-iT™ TMRM Reagent (I34361, invitrogen) according to the manual of manufacture. Fluorescent images were captured by Leica SP8 Confocal Microscope.

### Drug treatment

In ***Figure 1A-B***, Hela cells were treated with EBSS for 16 h, Torin1 (0.5 μM), CCCP (10 μM), rotenone (5 μM) for 24 h. In ***Figure S1A***, Hela cells were treated with Antimycin A (10 μM) and Oligomycin (10 μM), CCCP (20 μM) for 24 h. For cell viability in ***Figure 2A***, Hela cells were treated with mitochondrial toxin rotenone (10 μM) or CCCP (20 μM), ER stress inducer thapsigargin (TG, 1 μM) or lysosome inhibitor Brefeldin A (BFA, 1 μM) for 24 h. In K20-associated and ubiquitin associated experiments, cells were treated with CCCP (20 μM) for 6 h. For drug treatment in flies, flies were feed with food containing rotenone (250 μM), CCCP (100 μM) or thapsigargin (TG, 5 μM) for 7 days, and drug-containing fly food was changed every day. Flied was starved for 16 h with only water available before treatment.

### ATP Measurement

ATP was measured using an ATP Bioluminescence Assay Kit CLS II (11699709001; ROCHE) by following the manufacturer’s protocol and normalized with each thorax. The luciferase activity is measured on Lumat LB 9507 (Berthold Technologies).

### Statistical analysis

Data are expressed as mean ± SD. One-way ANOVA test with Tukey post hoc test and Student t-test were used for statistical evaluation. All statistical analyses were conducted using GraphPad Prism Software Version 9.0 (GraphPad Software Inc., La Jolla, CA). Cases in which P values of <0.05 were considered statistically significant.

## Acknowledgements

We are grateful to Drs. O. Brandman, E. Bennett, M. Hegde, S. Juszkiewicz for plasmids and antibodies; Drs. S. Birman, T. Littleton, J Chung, F. Gao, the Vienna *Drosophila* RNAi Center, FlyORF, and the Bloomington *Drosophila* Stock Center for fly stocks; Stanford PAN facility for primer synthesis; The Axelrod, Bogyo, Lipsick, and Svensson Labs in the Department of Pathology, Stanford University School of Medicine for sharing reagents and equipment. Special thanks go to J. Gaunce and W. Jiao for maintaining fly stocks and providing technical supports and members of the Lu lab for discussions. This work was supported by the NIH (R01NS084412 and R01AR074875 to B.L).

## Author Contributions

J.G designed the study, performed the experiments, analyzed data, and wrote the manuscript. Y.L, Z.W, and R.O performed experiments and analyzed data. S.L conceived and supervised the C9ALS part of the study, performed experiments, and provided funding. B.L conceived and supervised the study, performed experiments, wrote the manuscript, and provided funding.

## Additional information

### Supplementary figure legend

**Figure 1-figure supplement.**
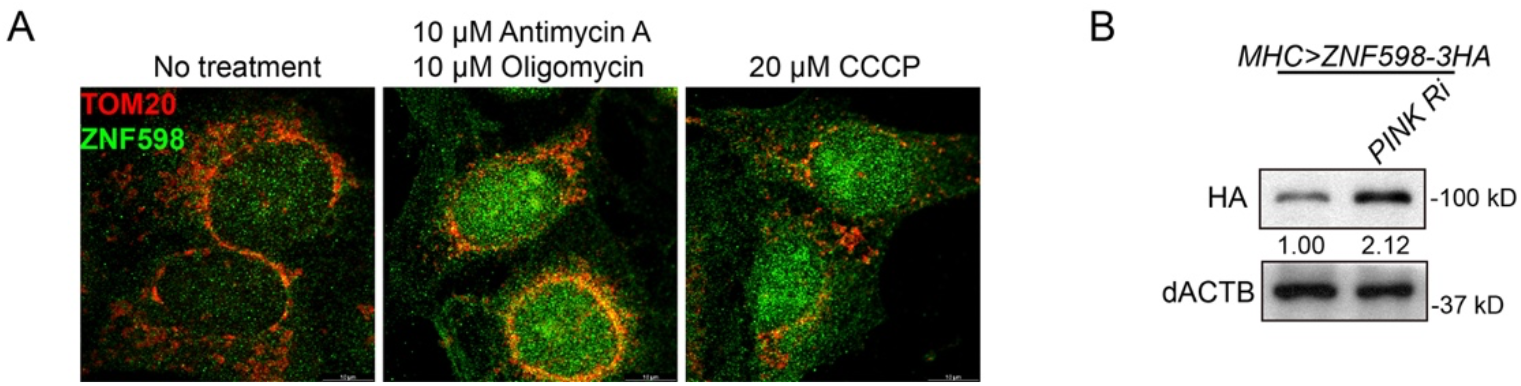
Upregulation of ZNF598 protein under mitochondrial stress *in vitro* and *in vivo*. (**A**) Immunostaining of ZNF598 (green) and the mitochondrial marker Tom20 in control cells and cells treated with CCCP or a combination of antimycin A and oligomycin. (**B**) Western blot analysis of HA-tagged ZNF598 transgene expression in *PINK1* RNAi fly muscle. All data are representative of at least 3 independent experiments. **Figure 1-figure supplement source data:** Western blots of HA-ZNF598 and Actin.

**Figure 2-figure supplement.**
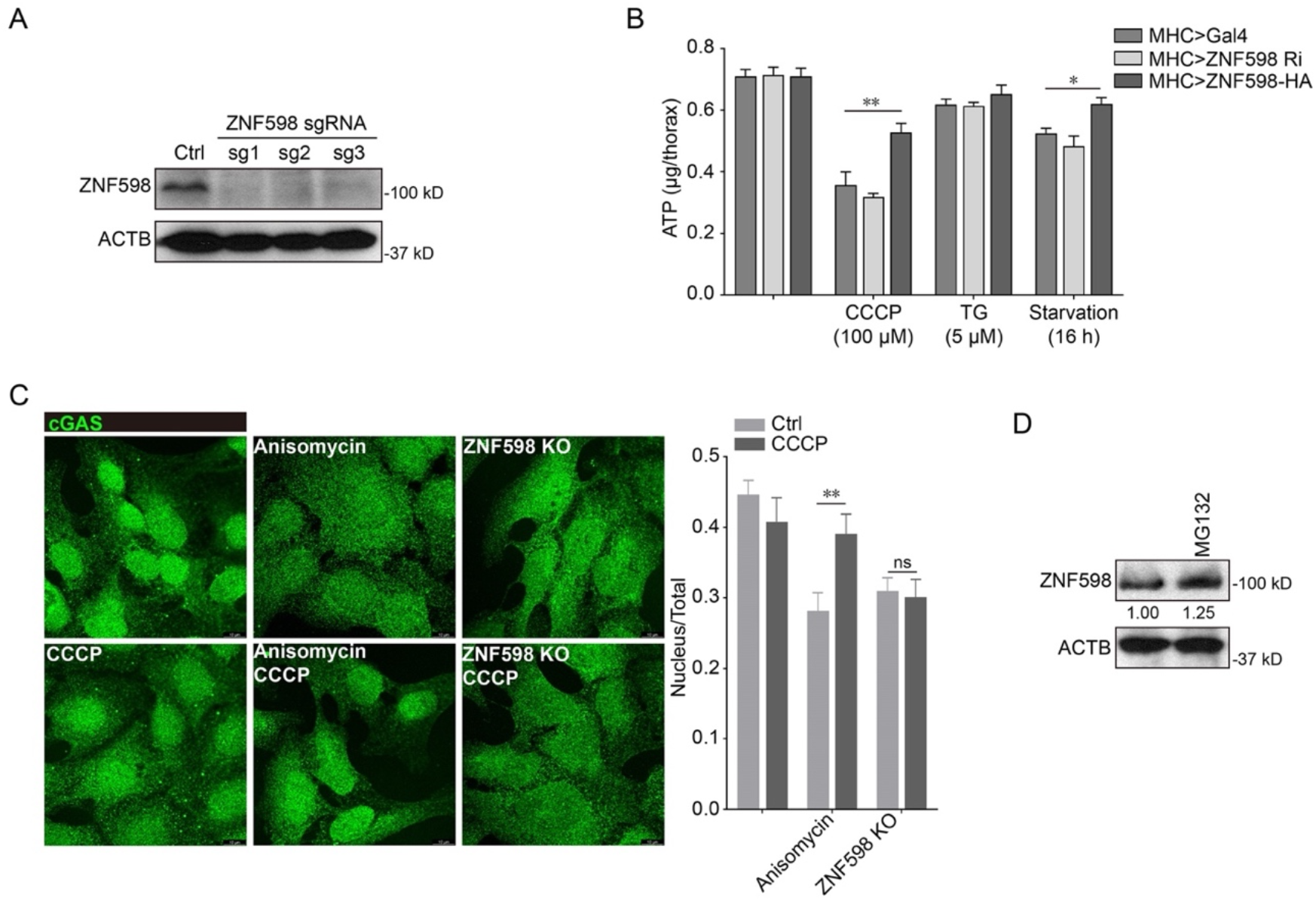
ZNF598 undergoes regulatory K63-linked ubiquitination under mitochondrial stress. (**A**) Immunoblots showing knockdown efficiency in CRISPR/Cas9-engineered ZNF598 knockout cells. (**B**) ATP measurement in control, ZNF598 overexpression, or ZNF598 RNAi flies after CCCP, TG, and starvation treatments for 16 h. n=3, *, P < 0.05 and **, P < 0.01 compared to *MHC*>*Gal4* control flies. (**C**) Immunostaining showing effect of anisomycin treatment or ZNF598 KO on cGAS subcellular distribution in U2OS cells with or without CCCP treatment. Bar graph shows data quantification. n=5, *, P < 0.05 and **, P < 0.01 compared with control. (**D**) Immunoblots showing effect of MG132 treatment on ZNF598 level in HeLa cells. All data are representative of at least 3 independent experiments. **Figure 2-figure supplement source data 1:** Quantification of ATP level and nuclear/cyto ratio of cGAS. **Figure 2-figure supplement source data 2:** Western blots of ZNF598 and Actin.

**Figure 3-figure supplement.**
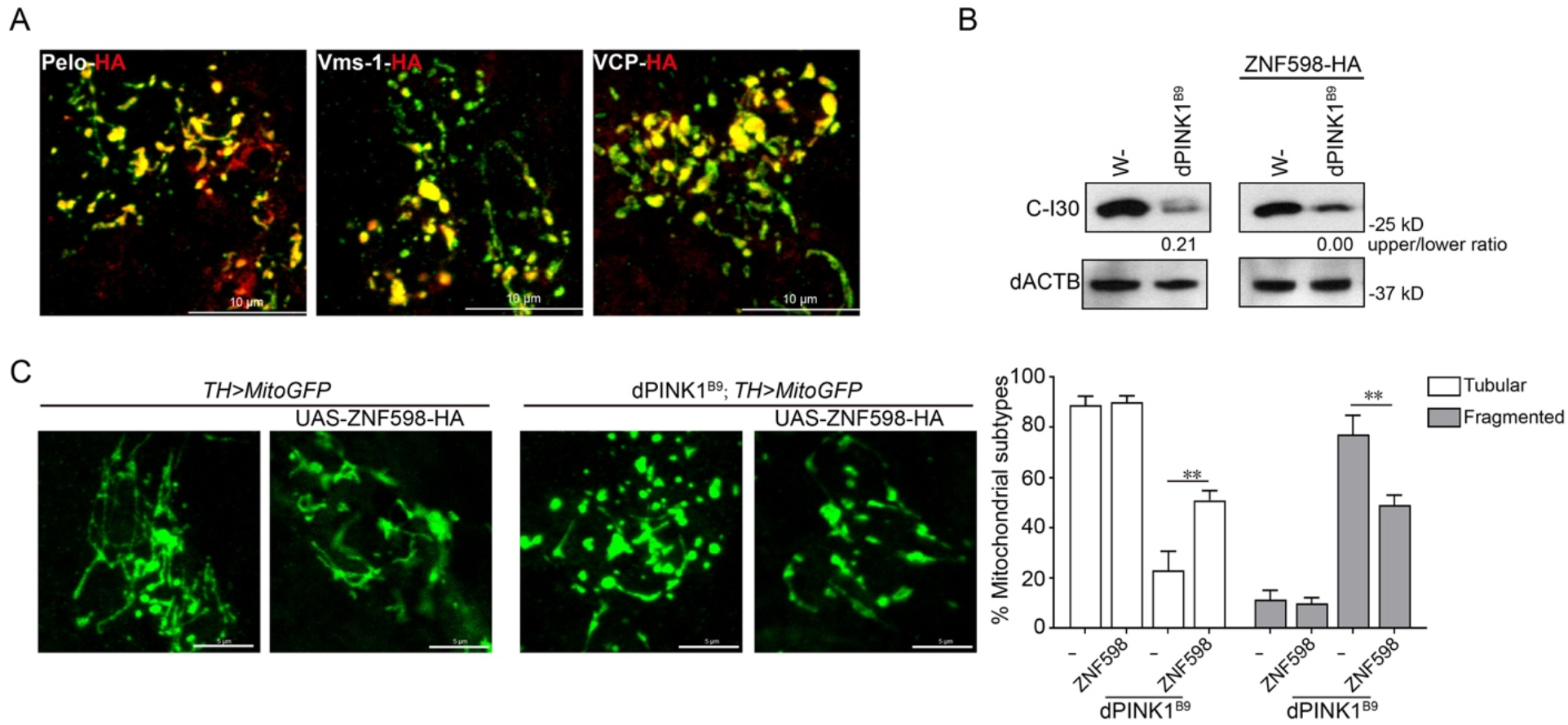
ZNF598 aborts stalled translation of mitochondrial outer membrane-associated *C-I30* mRNA and rescues mitochondrial and neuromuscular defects in *PINK1* mutant flies. (**A**) Localization of HA-tagged RQC factors Pelo, Vms-1, and VCP to mitochondria of adult fly DA neurons. DA neuron mitochondria are marked with *TH-Gal4*-driven mito-GFP expression. (**B**) Western blot analysis showing effect of ZNF598 overexpression on CAT-tailed C-I30-u formation in *PINK1^B9^* mutant flies. (**C**) Effect of ZNF598 overexpression on mitochondrial morphology in DA neurons of the PPL1 cluster of adult *PINK1^B9^* mutant fly brains. Bar graph shows quantification of relative proportions of tubular vs. fragmented mitochondria. All data are representative of at least 3 independent experiments. **Figure 3-figure supplement source data 1:** Quantification of mitochondrial morphology **Figure 3-figure supplement source data 2:** Western blots of C-I30 and Actin.

**Figure 4-figure supplement 4.**
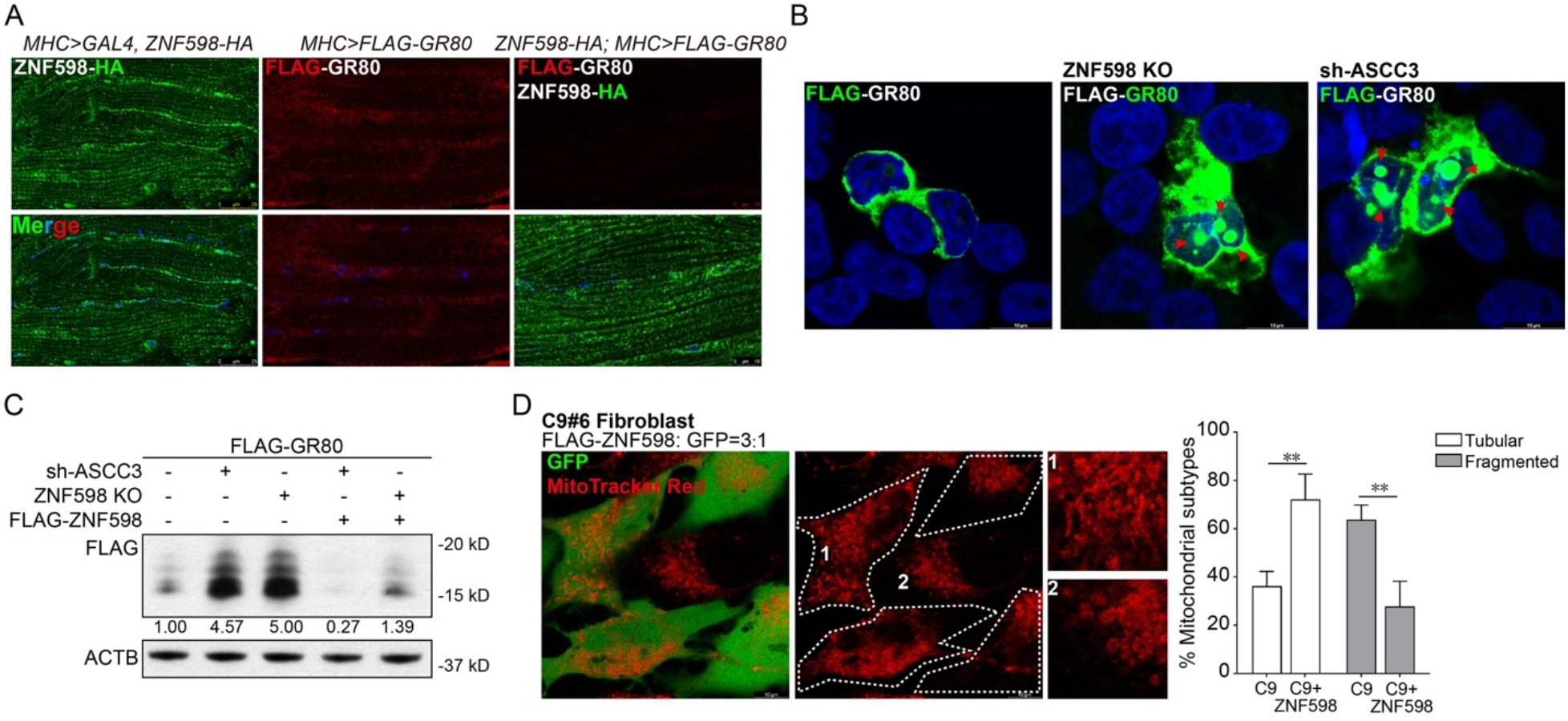
ZNF598 regulates the quality control of stalled translation of C9ALS-associated poly(GR) and rescues poly(GR) induced toxicity in *Drosophila* and patient cells. (**A**) Immunofluorescence staining for ZNF598 and FLAG-GR80 showing the effect of ZNF598 on GR80 protein level. (**B**) Immunofluorescence staining for FLAG-GR80 in ZNF598- or ASCC3-deficient HeLa cells. Arrowheads mark GR80 aggregates enriched in the nuclei of ZNF598- or ASCC3-deficient cells. (**C**) Immunoblots showing effect of ZNF598 overexpression on Flag-GR80 protein level in ZNF598- or ASCC3-deficient HeLa cells. (**D**) Mitotracker Red staining of ZNF598 transfected and non-transfected C9-ALS patient fibroblasts. Magnified views of cell 1 and 2 are shown on the right. Bar graph shows quantification of relative proportions of tubular vs. fragmented mitochondria. **Figure 4-figure supplement source data 1:** Quantification of mitochondrial morphology **Figure 4-figure supplement source data 2:** Western blots of Flag-GR80 and Actin.

